# Integration of silicon chip microstructures for in-line microbial cell lysis in soft microfluidics

**DOI:** 10.1101/2022.10.03.510547

**Authors:** Pavani Vamsi Krishna Nittala, Allison Hohreiter, Emilio Rosas Linhard, Ryan Dohn, Suryakant Mishra, Abhiteja Konda, Ralu Divan, Supratik Guha, Anindita Basu

## Abstract

The paper presents fabrication methodologies that integrate silicon components into soft microfluidic devices to perform microbial cell lysis for biological applications. The integration methodology consists of a silicon chip that is fabricated with microstructure arrays and embedded in a microfluidic device, which is driven by piezoelectric actuation to perform cell lysis by physically breaking microbial cell walls via micromechanical impaction. We present different silicon microarray geometries, their fabrication techniques, integration of said microarrays into microfluidic devices, device operation and testing on synthetic microbeads and microbial cells to evaluate their efficacy. The generalized strategy developed for silicon chip integration into soft polymeric devices can serve as an important process step for a new class of hybrid silicon-polymeric devices for future cellular processing applications. The proposed integration methodology can be scalable and integrated as an in-line cell lysis tool with existing microfluidics assays.

## I. INTRODUCTION

The use of soft microfluidic structures for cell sorting and genomic profiling is an active area of research and development, with powerful applications in medicine and biological research [1, 2]. While many microfluidic techniques have been successfully used to study mammalian single-cell genomics, similar studies of microbial cells have been partly limited due to the tough cell walls of many microbial species, leading to difficulty with in-line cell lysis, which is a critical first step needed to access the cellular contents within the microfluidic circuit. Partial success in single-cell microbial lysis had been achieved using lytic enzymes that target specific components in the cell walls [3, 4] or on microbes with lower cell wall thickness or strength, e.g., Gram-negative bacteria [5]. At present, there are no rapid, high-throughput lysis techniques that are compatible with microfluidics and which can be applied to a wide variety of microbial cells, including fungi, bacteria, etc. in an unbiased way. This can be particularly beneficial where *a priori* knowledge of the species present may be lacking, e.g., microbiome samples. We present a mechanical lysis approach to rupture microbial cell wall via micro-patterned silicon impactors integrated within PDMS-based soft microfluidic devices (**Fig. 1a**). This approach offers three advantages: 1) it is agnostic to the microbial species, 2) it is fast, lysing cells in less than a minute, and 3) unlike the enzymatic lysis approaches, it does not involve any chemical reagents that can potentially interfere with downstream experiments.

**Fig. 1.**
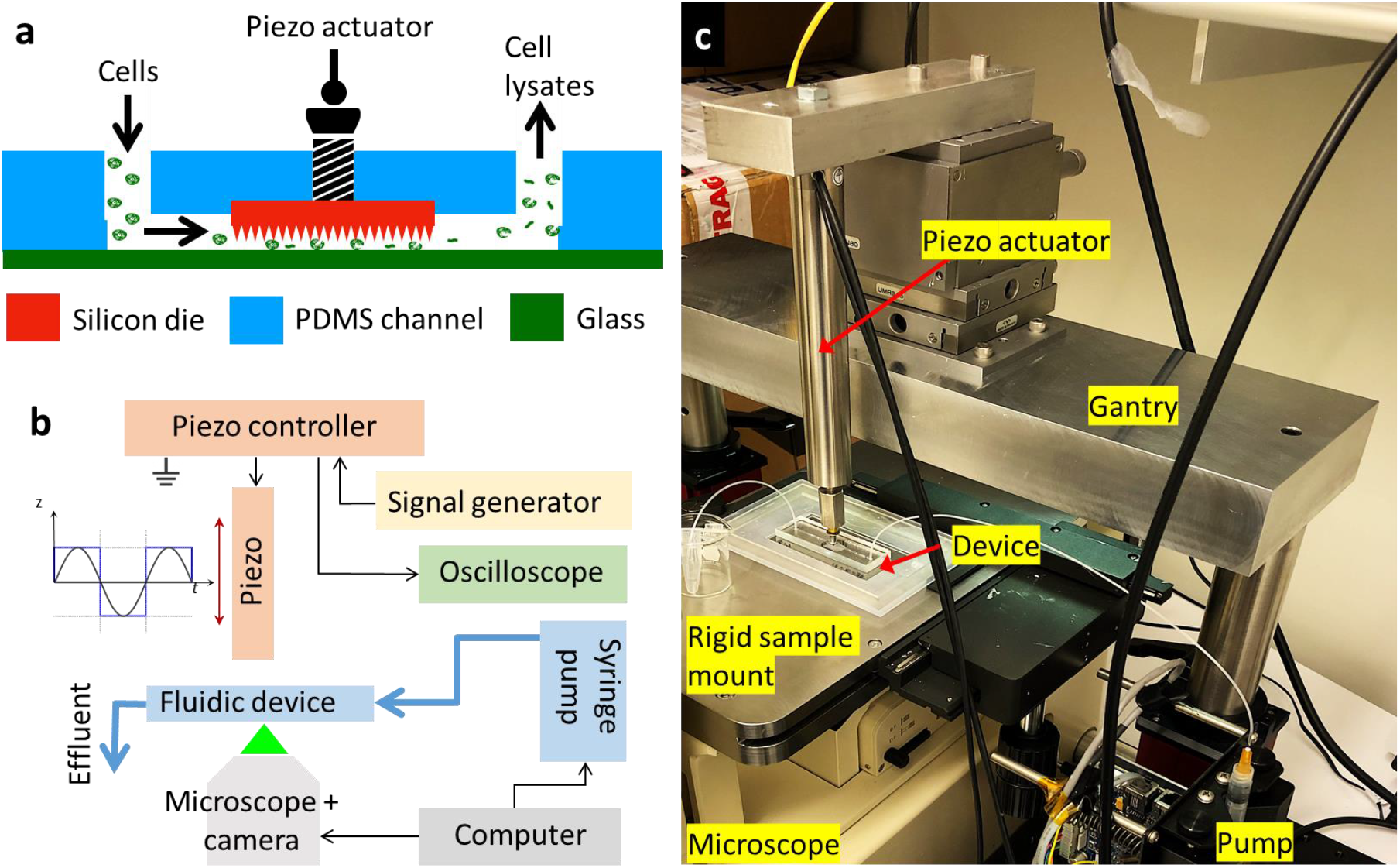
(a) Schematic showing the microfluidic device with embedded silicon chip and bonded against a glass substrate. A piezoelectric actuator will assist the silicon chip in moving up and down against the glass substrate to crush or lyse the cells. (b) Component diagram of the setup. (c) Custom setup showing the device on a microscope, connected to a pump and piezo actuator.

In this paper, we describe the process flow and fabrication of different patterned silicon micromechanical impactors developed for microbial cell lysis and their integration within the soft microfluidics environment. We developed an integrated architecture that combines mechanical drive motion of a patterned silicon chip for impaction, controlled flow in the microfluidic device, and optical imaging to visualize flow and mechanical impaction. Following initial testing and successful operation of the device (assessed on micron-sized silica and polystyrene beads suspended in water), we perform mechanical lysis of two microbial species, *Saccharomyces cerevisiae* and *Candida albicans*, using our approach. The current paper focuses mainly on the process development, fabrication and packaging of the device and its initial testing to demonstrate successful operation using both synthetic beads and microbial cells as test payloads.

We note that our integration approach is a general one for incorporation of semiconductor chip-based components into soft microfluidics that include developing process workflows that ensure mechanical flexibility of the device while providing leak-proof integration of the silicon chip into the soft polymeric device cast from a 3D printed mold. Such heterogeneous integration and packaging can open a range of new functionalities and capabilities within microfluidics applications well beyond lysis [6]. Optical, electrical, electrophoretic or RF interaction with single microbes may open up possibilities for additional sorting, manipulation and spectroscopic profiling [7]. With some modification, the current setup may also be employed for microbial single-cell ‘omics studies.

## II. EXPERIMENTAL SETUP AND METHODOLOGY

A schematic of the integrated package (**Fig. 1a**) and instrumentation (**Fig. 1b**) used for operation are shown. The patterned silicon impactor chip is embedded in the microfluidics device. An external syringe pump modulates the flow of cells within the microfluidic channels while piezo actuation drives the silicon chip back-and-forth, crushing the cells or beads against the glass substrate. Fabrication of different silicon chip designs and their integration into soft microfluidics are described as follows.

### A. Fabrication of Microarray-Patterned Silicon Impactor Chips

Microstructural features were patterned using optical lithography followed by dry or wet etching. All 4-inch Si wafers (Silicon Valley Microelectronics, Cat# SV007) were quartered, sonicated in acetone and isopropanol for 3 min each, and then rinsed in water and dried at 115 °C on a hot plate for 1 min prior to further handling. Patterns used for optical lithography were designed using Siemens L-Edit software and defined via optical lithography using the Heidelberg MLA150 maskless aligner (at 405 nm), with varying photon doses as outlined below. Descum was performed prior to etching and deposition steps using a CS-1701 Nordson March etch tool (Westlake, OH) set to 50 W, 160 mTorr chamber pressure, and with an O^2^ flow of 24 standard cubic centimeters per minute (sccm) for varying amounts of time detailed later where needed. Chromium deposition (with typical layer thicknesses of 20 nm) was carried out using a Temescal FC-2000 E-beam Evaporator. Deep reactive ion etching (DRIE), BOSCH-like etching [8], and reactive ion etching (RIE) were performed on an Oxford PlasmaLab System 100. In addition, DRIE etching was also carried out also using a DRIE tool from PlasmaTherm following the standard Versaline process. Five different patterned silicon microarrays were fabricated using the process flows outlined:

#### KOH Pyramids

A Si <100> wafer with 1000 nm thick thermal silicon dioxide (Si/SiO_2_) substrate (**Fig. 2a**) was spin coated with 1.3 µm thick positive resist (Shipley S1813, Marlborough, MA), and baked at 115 °C for 1 min. The resist was then patterned into 2, 3, and 4 µm squares with 1 µm spacing between them (**Supplementary Fig. S1a**). The resist was developed in 1:3 v/v developer AZ 351 (MicroChemicals GmbH, Germany): deionized (DI) water for 22 sec, rinsed in DI water and hard-baked for 3 min at 115 °C. The sample was treated with a weak O_2_ plasma (descum) for 30 sec to remove any resist residue in the developed regions after the lithography (**Fig. 2b**) and then transferred to the RIE system to etch the Si/SiO_2_. For this, a 500 W Inductively Coupled Plasma (ICP) was used with the lower electrode (platen) at 25 W, chamber process pressure at 20 mTorr and the electrode temperature of 20 °C (**Fig. 2c**). The etch time was typically 30 min with an O_2_ flow of 1.0 sccm and a CHF_3_ flow of 55 sccm. The resist mask was then removed in Microposit resist remover 1165 (Dupont, Wilmington, DE) via a 6 hr immersion. The remaining structures are then wet etched anisotropically in a 30% KOH solution at 90 °C [9-12], producing square-based pyramids (**Fig. 2d, o**) via the process of undercutting beneath the oxide layer.

**Fig. 2.**
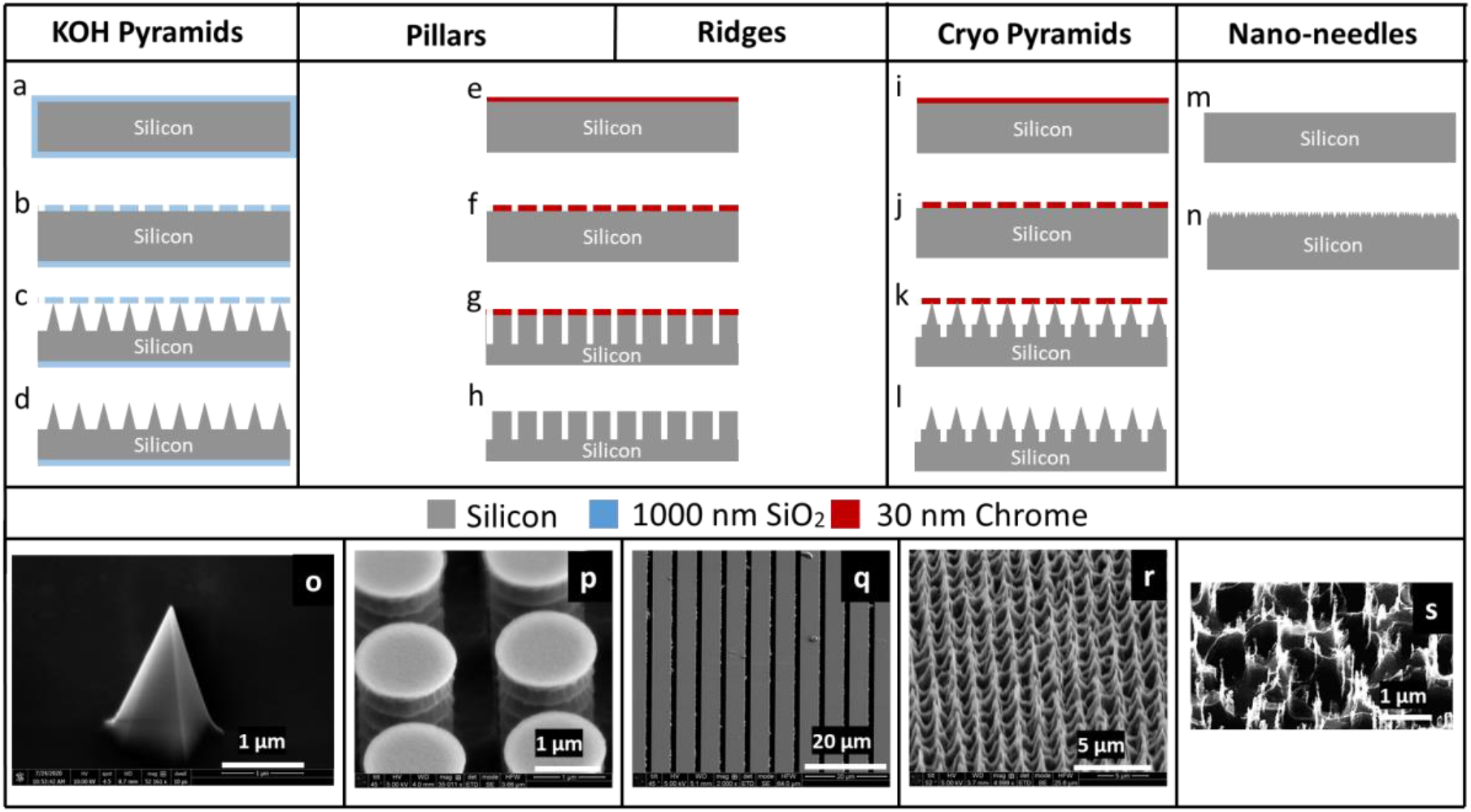
Silicon chip fabrication. Process flow to fabricate: (a-d) pointed pyramidal structures, using the KOH-based etching, (e-h) high-density pillars or ridges, using BOSCH etch in DRIE; the same process flow yields two different geometries when using different lithography masks and etch times, (i-j) high-density cryo pyramids, using cryo-etch process, (m-n) nano-needles, using DRIE. SEM images showing (o) a single pyramid fabricated using KOH etching of Si; (p) Array of dense cylindrical structures fabricated using BOSCH etch process; (q) Array of ridges fabricated using BOSCH etch process; (r) Array of dense cryo pyramids using cryo-etch process; (s) Array of nano-needles with features of 15-20 nm fabricated using BOSCH etch process.

#### Pillars and Ridges

A Si <100> wafer was spin coated with 270 nm thick positive resist S1805 (Shipley): propylene glycol methyl ether acetate (PGMEA, Sigma, Cat# 484431) at 2:1 v/v ratio and baked at 115 °C for 1 min (**Fig. 2e**). The resist was then patterned with either 1.1 µm circles with 0.9 µm gaps, or grids with a width of 5 µm and a gap of 1 µm to generate to generate pillars or ridges, respectively (**Supplementary Fig. S1b-c**). The pattern was developed in 1:4 v/v Developer AZ 351: DI water for 22 sec and the residual resist was descummed for 15 sec. After this, the wafer was coated with Cr and after lift-off with resist remover 1165 for 6 hr to overnight, the Cr hard mask was obtained (**Fig. 2f**). This pattern was transferred to the Si wafer by a BOSCH etch process. This process consists of a sequence of repeated alterations between deposition and etch steps with C_4_F_8_ and SF_6_, respectively [8] (**Fig. 2g**). The deposition step was performed at 700 W ICP power, 10 W lower electrode (platen) and with a C_4_F_8_ /SF_6_ gas flow of 100 sccm/1 sccm. The etch step was performed at 700 W ICP power, 30 W platen power, with C_4_F_8_ /SF_6_ gas flows of 1 sccm/80 sccm. During the deposition and etch steps, the chamber process pressure was 30 mTorr and the electrode temperature was held at 0 °C. To generate pillars (**Fig. 2h, p**), this process was repeated for 12 cycles; to generate ridges (**Fig. 2h, q**), this process was repeated for 50 cycles, where one etch step and deposition step together are considered a full cycle.

#### Cryo Pyramids

A Si <100> wafer was spin coated with 270 nm thick S1805: PGMEA and baked at 115 °C for 1 min, as described above (**Fig. 2i**). The resist was patterned (same as **Supplementary Fig. S1b**) and then developed in 1:4 v/v Developer AZ 351: DI water for 22 sec. Following descum process for 15 sec, the wafer was coated with a Cr hard mask as described above (**Fig. 2j**). Finally, the pattern was transferred to Si wafer (**Fig. 2k**) by a low temperature (−90 °C) cryo based silicon RIE process [13, 14], producing sharpened pyramidal structures (**Fig. l, r**) by lateral undercutting of the Cr mask, with process conditions 700 W ICP power, 3 W Platen power, chamber pressure of 5 mTorr, the electrode temperature at -90 °C and with O_2_ flow of 4.0 sccm, CHF_3_ flow of 6 sccm, SF_6_ flow of 32 sccm, and etch rates ∼350 nm/min.

#### Black silicon nano-needles

A Si <100> wafer was quartered, cleaned, dried (**Fig. 2m**), and etched directly. To ensure the plasma is sustained during the etch process, a brief strike step with the following conditions is introduced: 1500 W ICP power, chamber pressure was 10 mTorr, the electrode temperature was 15 °C for a duration of 5 sec with Ar flow of 30 sccm and C_4_F_8_ flow at 75 sccm. After this, a 5 min etch was carried out with 1200 W ICP power, chamber pressure was 20 mTorr, the electrode temperature was 15 °C for 5 min with O_2_ flow of 50 sccm and SF_6_ flow at 70 sccm to generate an array of disordered black silicon (bSi) nano-needles, consisting of nanoscale features with a high aspect ratio (**Fig. 2n, s**). The BSi nano-needles have demonstrated mechanical bactericidal effect [15, 16].

After the microarray patterns were fabricated, the Si wafer quarters were coated in resist S1813 at 1.5 µm thickness and diced into 5×5 mm_2_ chips, using an ADT 7122 dicing tool. Any residual resist was stripped in resist remover 1165, and the chips were rinsed in DI water and dried. At this point, the silicon chips were ready to be embedded into the microfluidic device.

### B. Microfluidic Device Fabrication and Silicon Chip Integration

Assembled devices are composed of a soft elastomeric polydimethylsiloxane (PDMS) block with a microfluidic channel patterned via soft lithography, a microarray-patterned silicon chip inset into the PDMS block, a glass substrate, and a metal screw (**Fig. 1a**).

**Fig. 3** illustrates the schematics of the process flow to embed the silicon impactor chip into the soft microfluidic device. Devices were designed and assembled using standard soft lithography [17] processes. Briefly, a PDMS block was generated by replica molding around a resin mold patterned with the inverse of a microfluidic channel, and a square inset of 5×5 mm_2_ for seamless inlay of the microarray-patterned silicon chip. The resin master mold was designed in AutoCAD (AutoDESK, USA) (**Supplementary Fig. S1d**) and printed using a Form 3 3D printer (Formlabs, Somerville, MA) outfitted with V4 Clear Resin (Cat# RS-F2-GPCL-04) set to a process resolution of 25 µm. The printed material was soaked in an IPA tank for 3 hr and UV cured at 75 °C for 7 hr to generate the mold (**Fig. 3j**) patterned with the inverse of features desired in the final PDMS block, including raised portions which later define the microfluidic channel and an inset for the silicon chip. Typical channel dimensions in the microfluidic device are 125 µm deep, 1500 µm wide with a recess of 5000×5000×450 µm_3_ to fit the silicon impactor chip. Since the actual thickness of the silicon chip is ∼503-504 µm, there is an overhang of 53-54 µm for the silicon chip into the flow channel (**Fig. 3i**, not drawn to scale). To cast the mold, PDMS elastomer and cross-linker (Krayden Dow Sylgard 184 Silicone Elastometer Kit) were mixed at a 1:10 w/w ratio using a THINKY AR-100 centrifugal mixer (Laguna Hills, CA) set to 30 sec mixing and 30 sec degassing, then poured into the resin mold (**Fig. 3a-b**), degassed gently under vacuum for 20 min to remove air bubbles, and subsequently baked in an oven at 65 °C for >3 hr to cure the PDMS. The cured PDMS was then cut away from the edges of the mold using a scalpel and peeled off, leaving a patterned PDMS block (**Fig. 3c**). Subsequently, 4 mm sized biopsy punch (Robbins Instruments, Cat# RBP-40) was used to cut a hole at the center of the silicon die inset (**Fig. 3d**) and a 1 mm biopsy punch (Integra Miltex, Cat# MIL-33-31AA-P/25) was used to cut inlet and outlet holes to the microchannel (**Fig. 3e**). The PDMS block was then cleaned by sonication in DI water for 3 min and IPA for 6 min to prepare for incorporation of the silicon chip. The silicon chip was inlayed and bonded to the elastomer surface by applying a minute amount of uncured PDMS elastomer and cross-linker mix, prepared as described above, to the PDMS at the corners of the inset and then gently pressing the silicon chip, microarray-patterned side facing outward, into place until flush with the microchannel. The patterned PDMS block and silicon chip were then baked, chip-side down, on a 95 °C hotplate for 30 min (**Fig. 3f**) to cure the fresh PDMS. Following this, the microfluidic device with embedded silicon chip is ready to be sealed.

**Fig. 3.**
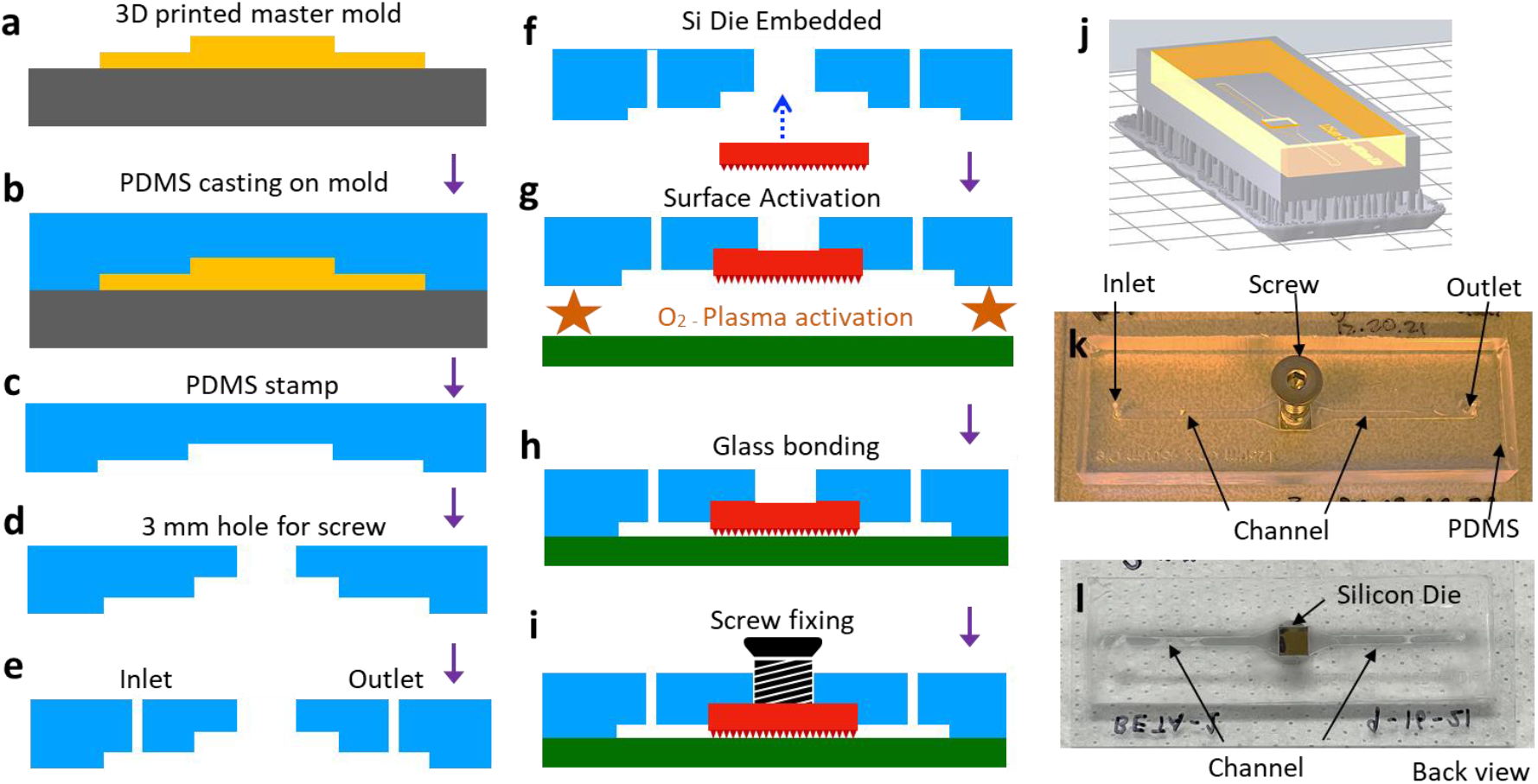
Schematic for device integration. (a) 3D printing the master mold, (b-c) Fabrication of PDMS stamp from master mold, (d-e) Punching the 3 mm hole and the fluid inlet/outlets using biopsy punch, (f) Embedding patterned silicon chip and bonding using uncured PDMS, (g-h) Oxygen plasma treatment and bonding of glass slide and PDMS stamp, (i) Inserting a M3 screw and sealing the opening using permanent epoxy, (j) CAD schematic with raft (yellow) and support (gray), (k-l) Images of the front and back views of the device showing screw, die and inlet, outlet of the channel.

Glass slides 75×38 mm_2_ (Fisher Scientific, Cat# 12-550B) were cleaned by scrubbing in acetone, then sonicating in acetone for 30 min, IPA (>99.5%) for 10 min, and DI water for 10 min. A 5×5 mm_2_ Si <100> piece was placed over the patterned chip to protect its microstructures, while a BD-20AC Laboratory Corona Treater (Electro-technic Products, USA) was used to apply air plasma alternatingly to the PDMS surface and the cleaned glass slide for a total of 10 min. After this, the protective Si piece was removed and the treated surfaces were pressed together to create a covalently bonded, leak-proof microfluidic channel. The patterned silicon chip remained suspended over the microfluidic channel (**Fig. 1a**). The device was then baked on a hot plate at 75 °C for 30 min (**Fig. 3g, h**) to strengthen the covalent bonding between the PDMS block and glass slide. Finally, a M3 screw was inserted into the 3 mm hole behind the silicon chip and attached with a small amount of Devcon 2 Ton Epoxy (**Fig. 3i**), providing a rigid external contact point through which mechanical motion can be transmitted to the embedded chip. The completed device (**Fig. 3k, l**) is tested for leakage by flowing DI water in the microchannel before it is deemed ready for use.

### C. Cell Culture and Fluid Sample Preparation

We performed preliminary impaction tests with porous polystyrene-divinylbenzene beads of 10 µm diameter (Chemgenes) and porous silica (SiO_2_) beads of 3 µm diameter (Sigma, Cat# 806765). The 10 µm beads came functionalized with hexaethylene glycol and DNA fragments that prevented aggregation. Both 10 µm and 3 µm beads were suspended in DI water at 4.5 million and 65 million beads//mL, respectively. The beads were loaded into syringes (BD biosciences, Cat# 309657) for flow experiments.

Next, we used fungal microbes to test lysis from impaction experiments. Two species, *Saccharomyces cerevisiae* and *Candida albicans* were tested separately for comparison. *S. cerevisiae* (strain BY4741, Open Biosystems) and *C. albicans* (strain SC5314, ATCC) were grown overnight in YEPD medium (MP Biomedicals, Cat# MP114001022) and used in log phase. To prepare the cells for impaction experiments, cells from each species were spun down in a centrifuge at 130 xg for 4 min and the supernatant medium was removed. The cells were washed and spun down at 130 xg for 4 minutes in 1X PBS (Fisher BioReagents 10X PBS, Cat# BP399500, diluted to 1X using Invitrogen Ultrapure DNase/RNase

Free Distilled Water, Cat# 10977023), and re-suspended in 1X PBS. 10 µL of the suspension was removed from the stock and counted on a hemocytometer (InCyto, Cat# DHC-N01-2) and the cell concentration was adjusted to generate 1 mL of cell suspension in 1X PBS at 50 million cells/mL and 400 U/mL RNase Inhibitor (Lucigen, Cat# F83923). The cells were stored on ice and loaded into sterile plastic syringes (BD biosciences, Cat# 309657) immediately before use. Effluents were collected on ice to preserve RNA integrity.

### D. Device Operation and Sample Collection

During the silicon chip integrated PDMS device operation, the inlet and outlet channel are used to flow solutions that contain the test specimens (e.g., microbeads or microbes). The patterned silicon chip is used as an impactor to mechanically squish or crack the microbeads or microbial cells under flow. When the piezo actuator is activated at an applied frequency and waveform, the silicon chip is pushed by the actuator to move up and down continuously such that the microbes or test specimens underneath are squished between the Si structures and the glass substrate (**Fig. 1a**), with the glass substrate acting as an anvil. This piezoelectricity-driven micromechanical actuation of the different fabricated microstructures on the silicon impactor chips (pointed pyramids, pillars, ridges, etc.) causes the microbeads or microbial cell walls to either perforate, crack or break due to applied stress, thereby lysing the cell.

Vertical motion of the silicon chip was generated using a piezoelectric actuator (P-841.40 piezo actuator model; E-665.SR Piezo amplifier/servo controller) (**Fig. 1a, c**). A square waveform with peak-to-peak voltage (V_pp_) = 10 V, duty cycle = 50%, and frequency = 0.5 Hz was applied to the piezo amplifier using a BK Precision 4053B waveform generator. This translated into linear displacements of ∼40 µm by the piezoelectric element, which coupled to the silicon chip embedded in the fluidic device through the M3 screw (**Fig. 1a; 3i, k**), resulting in linear displacement of the silicon chip into the microfluidic channel and its contents in the same frequency-dependent manner that affects crushing (or puncturing) of the payload in the channel (e.g., microbial cells or micron sized beads ∼1 - 10 µm in diameter, used as proxy for microbes).

Experiments were carried out on a Nikon Epiphot 200 inverted metallurgical microscope, equipped with extra-long working distance 20x and 40x air objectives and a MiChrome 5 Pro (Tucsen) camera to assist with alignment of the piezoelectric element, monitor sample flow through the device and assess silicon chip displacement (via change in focal plane). A custom-built rigid baseplate (**Fig. 1c**) was installed as the sample holder to minimize sag/deformation of the device under mechanical impaction. Likewise, the piezo element was mounted on a mechanically rigid steel gantry (**Fig. 1c**) to allow the full force of the piezoelectric motion transmitted to the silicon chip impaction. The gantry design also incorporates an X-Y-Z differential translation stage (Thorlabs, Cat# PT3A) to allow easy alignment of piezo element with the fluidic device while visualizing flow and mechanical impaction.

The silicon impactor chip integrated microfluidic device was placed on the baseplate such that the patterned impactor surface could be observed in the microscope’s field of view. The sample of interest (either micro-beads or live cells suspended in fluid) was flowed through the device at 300 µL/hr using a KDS 910 syringe pump (KD Scientific) and controlled using custom code written in LabVIEW (National Instruments) until the microchannel was completely filled with liquid, displacing any trapped air pockets). Undisturbed flow was maintained for 10 min, discarding the effluent. 50 µL of effluent (without piezo-electrically driven impaction) was collected for ∼10 min as negative control.

Next, the flow was turned off to allow proper alignment of the piezoelectric head. The piezoelectric head was aligned and brought into contact with the center of the flat top of the M3 screw, with contact indicated by the observation of a slight change in microscope focus corresponding to a change in the z-position of the chip (induced by a small change in the local channel width as it deformed under mechanical loading).

Following alignment, flow was resumed through the device at 300 µL/hr and piezoelectric actuation in the form of an oscillatory square wave of V_pp_ = 10 V was applied. After waiting an additional 10 min for residual volume not brought into contact with the actuated silicon chip to flow through and be discarded, 50 µL of the effluent was collected directly into a 200 µL pipette tip with barrier filter (Genesee Scientific, Cat# 24-412) and saved for analysis.

### E. Scanning Electron Microscopy: Sample Preparation and Imaging

We evaluated the system’s ability to mechanically crush microbeads and microbial cells using scanning electron microscopy (SEM). We performed SEM on bead and *S. cerevisiae* cell residues on the surfaces of silicon chip impactors retrieved after the experiments, and also in the effluent fluid collected from the outlet of the fluidic device.

Effluent samples were prepared for SEM imaging by placing a small droplet of effluent on a plain Si <100> 5×5 mm_2_ chip and allowing it to dry overnight (or longer). *S. cerevisiae* effluent samples were prepared for SEM imaging following a modified version of the procedure described in [18]. Briefly, glutaraldehyde (GLA; Electron Microscopy Sciences, Cat# 16120) was added to the effluent to a final concentration of 2.5% GLA and the samples were left to fix in a 4 °C fridge for 2 hrs. Following fixation, cellular samples were spun down and washed once with 1X PBS at 130 xg for 4 minutes to remove excess glutaraldehyde and then dehydrated through a series of ethyl alcohol washes at 30%, 50%, 70%, and twice at 100%. A small droplet of the dehydrated sample was placed on a 5×5 mm_2_ piece of plain Si and allowed to dry overnight.

Samples of crushed microbeads and *S. cerevisiae* cells on the patterned silicon impactor chips were prepared by allowing the microfluidic device to dry overnight and then extracting the silicon chip from the microfluidic device using a scalpel. The silicon chip was further dried at room temperature overnight. The dried samples were coated with a thin layer of gold (a few nm thick) by sputter coating (Cressington Sputter Coater 108 Auto, Ted Pella, Inc.) for 120 sec to enhance contrast under SEM. SEM images were collected using a FEI Nova NanoLab DualBeam focused ion beam (FIB) instrument equipped with high resolution electron beam for SEM.

### F. SEM Image Analysis to Characterize Crushed Microbeads

The images of effluent samples were sorted according to the size of the microbeads (3, 10 μm) and the type of silicon microarray pattern used to crush them. Multiple SEM images of 10 μm (acquired at ∼650x magnification) and 3 μm (2000x magnification) microbead samples were used to calculate crushing efficiency.

The SEM images were analyzed using the cell counter plugin in NIH ImageJ software. Crushing efficiency was calculated as the number of crushed beads divided by the total number of beads (crushed and uncrushed). The number of crushed beads was calculated by estimating the size of visible crushed fragments and multiplying their approximate size by the number of fragments of that size. For simplicity, all visible bead fragments were binned as follows: intact, ½, ¼, ⅛, 1/16, 1/32 and 1/64 times the bead size. Full beads were counted first, and then crushed bead fragments were counted by their approximate sizes. Intact beads on the borders of the image were included. A tally of the number of beads versus the total number of crushed beads, calculated by multiplying the total count for each size of fragment by the approximate size of the bead fraction and summing the counts from all fractions, was recorded. The error bars represent the standard error according to data taken from multiple images.

### G. qRT-PCR Based Quantification of Yeast Cell Lysis on Chip

Real-time quantitative reverse transcription-polymerase chain reaction (qRT-PCR) was used to quantify the RNA released into the effluent medium following impaction experiments on *S. cerevisiae* and *C. albicans*. qRT-PCR was performed on the effluent lysate at different dilutions, along with positive and negative controls, using QuantStudio 3 Real-Time PCR, 96-well, 0.2 mL System (Applied Biosystems) and iTaq Universal SYBR Green One-Step Kit (BioRad, Cat# 1725150). PCR primers for *S. cerevisiae* [19] and *C. albicans* [20] were designed and custom-synthesized from IDT.

We used qRT-PCR to detect the presence of mRNA for three *S. cerevisiae* genes: *ACT1, UBC6* and *TDH3*. These genes are good candidates to assess cell lysis due to their high (*TDH3, ACT1*) or moderate (*UBC6*) expression levels [19]. Crushed cell lysate (indicated as ‘Sample’) was collected as effluent after flowing through the device under piezo actuation and tested at stock concentration and dilutions of 1:10 and 1:100 (denoted as ‘Sample 1:10’ and ‘Sample 1:100’). Effluent of uncrushed *S. cerevisiae* cells was also collected under flow but with the piezo actuation turned off to be used as a negative control effluent (indicated as ‘N. Control E’). Nuclease-free water was tested as an independent negative control (‘N. Control’). As positive controls (‘P.C.’) for qPCR, we used 50 million *S. cerevisiae* cells lysed using zymolyase and sarkosyl [21], followed by RNA purification with an extraction column (Zymo Research, Cat# R2070) and elution in 100 µL. Stock solution of P.C., along with dilutions of 10x, 100x and 1000x were used for *TDH3, ACT1*, and *UBC6* in *S. cerevisiae* experiments.

We also used qRT-PCR to detect mRNA for three *C. albicans* genes: *LSC2, TDH3* and *ACT1. TDH3* is one of the highest expressed genes in *C. albicans* [21]. Crushed cells collected as effluent after flowing through the device under piezo actuation were tested at stock concentration (indicated as ‘Sample’) and dilutions of 10x and 100x (denoted as ‘Sample 1:10’ and ‘Sample 1:100’). Uncrushed *C. albicans* cells were also collected under flow but with the piezo actuation turned off as the negative control effluent (indicated as ‘N. Control E’). As before, we used 50 million *C. albicans* cells lysed using zymolyase and sarkosyl [21], followed by RNA purification and elution in 100 µL. Stock solution of the lysate denoted as ‘P.C.’, along with dilutions at 5x, 10x, 20x and 100x were used for *LSC2, TDH3* and *ACT1*. PCR amplification experiments were run for 40 cycles, where we see late amplification for negative controls (all control data are not shown).

## III. RESULTS AND DISCUSSION

### A. Silicon Microarray-Patterned Chip Fabrication

SEM images of the nanofabricated microarray patterns show the successful generation of several different types of structures (**Fig. 2**). KOH Pyramids: 2 µm wide, ∼2 µm high, 4-5 µm spacing; Pillars: 1.1 µm wide, 5 µm deep, 0.9 µm spacing; Ridges: 5 µm wide, 40-50 µm deep, 1 µm spacing; Cryo Pyramids: 1.5 µm wide, ∼2-3 µm high, 1 µm spacing; Nano-needles: 50-200 nm wide, 0.5-1 µm high, 0.3 µm – 0.4 µm spacing.

### B. Analysis of Micro-bead Impaction via SEM Image Analysis

SEM images show the effect of impaction upon the beads using different silicon chip patterns (**Fig. 4a-o**). We observed bead breakage following impaction in all cases, but the percentage of crushed beads depended on silicon chip geometry and beads used. In some cases, we observed instances where the 3 µm beads were perforated by sharp KOH pyramid tips (**Fig. 4a**). Some beads were also crushed (**Supplementary Fig. S2a**) or ripped away (**Supplementary Fig. S2b**) by the KOH pyramid tips. 3 µm beads were either crushed or embedded in the pillar microarray (**Fig. 4d, e, Supplementary S2c, d**). We also saw cases where the 3 µm porous silica beads distorted the Si pillars and damaged them. To bypass this, we employed the ridge geometry (**Fig. 4g, h**), which is mechanically stable and provides improved strength to the silicon impactor chip. We also found that fewer beads were crushed when the Si ridges were perpendicular to the direction of the flow, instead of being aligned with flow (**Supplementary Fig. S2e, f**). When the channels between the ridges are aligned with the flow, the fluid squeezed into the channels by the impaction process can escape forward and backward into the microfluidic channel. When the channel is perpendicular to the flow, this is not possible and a back pressure develops opposing the impaction motion. The ridged silicon chips have crushing efficiencies between 10.3 – 32.8%, depending upon size of the beads.

**Fig. 4.**
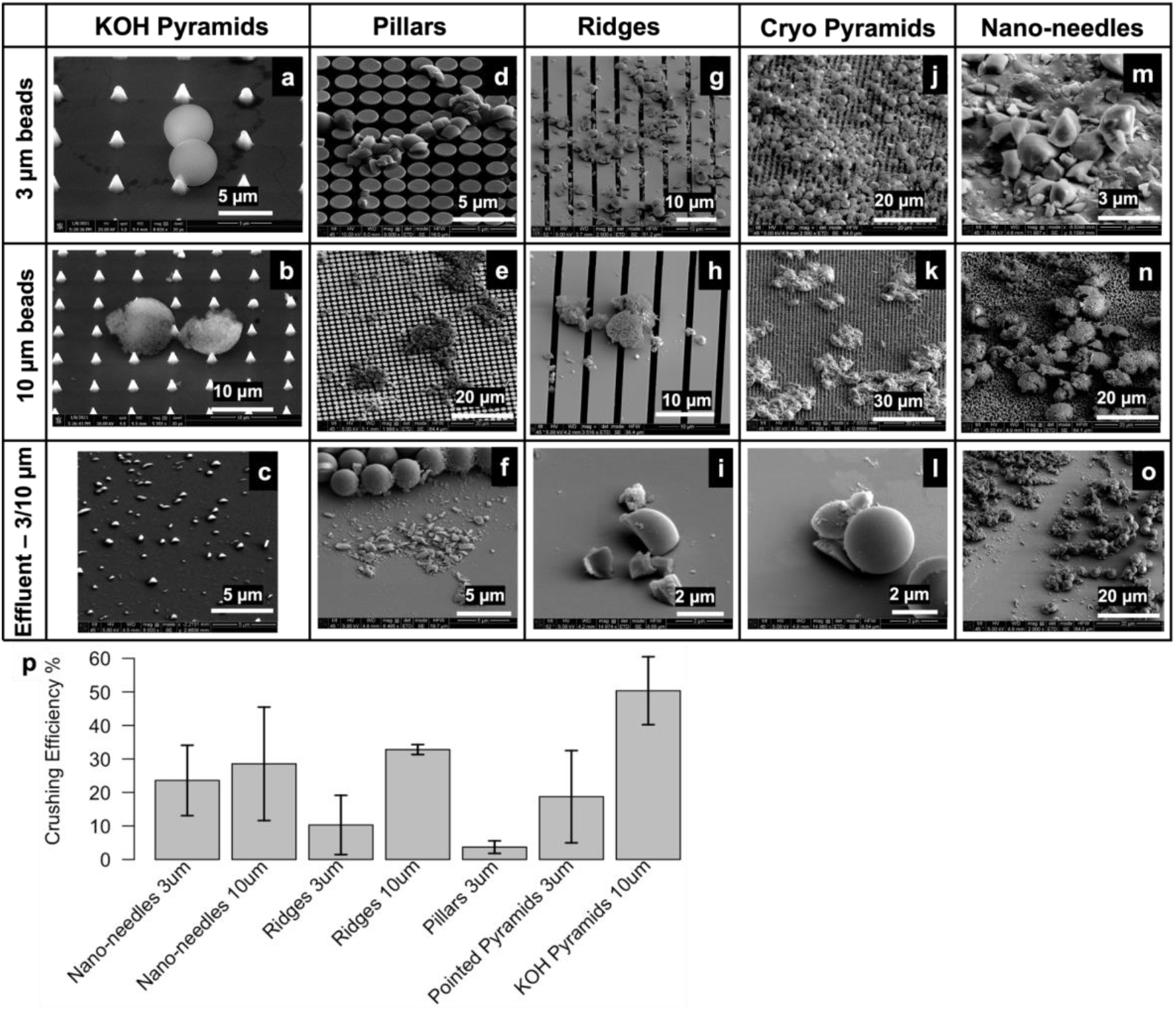
SEM images showing the damage to top- 3 µm beads on chip, middle- 10 µm beads on chip, or bottom- effluent (3/10 µm beads), caused by silicon chip variant (a-c) KOH pyramids, (d-f) Pillars, (g-i) Ridges, (j-l) cryo pyramids, and (m-o) Nano-needles. (p) Bead crushing efficiency reported as the percentage of beads crushed vs. silicon impactor chip type and bead size.

Detailed analyses of SEM images (described above) of the effluent collected on Si wafers were performed to estimate the percentage of crushed beads. The crushing efficiencies for different silicon chip types and bead sizes are compared (**Fig. 4p**). Overall, microbead crushing efficiencies varied between 3.7 – 50.4 % for different silicon chip geometries. For any given silicon chip type, we noted higher crushing efficiency in 10 µm beads compared to 3 µm beads, as the probability of the 10 µm beads escaping between the silicon structures is lower. For example, the spacing between the KOH pyramids is ∼4-5 µm, which the 3 µm beads can escape into, without being crushed. Reducing the gap between the KOH pyramids is difficult due to the crystallographic orientation of Si used in the etching process. The KOH pyramids showed the maximum crushing efficiency (∼50.4 ± 5.1 %) using 10 µm beads.

These results demonstrate that the silicon chip travels to within 10-3 µm from the glass substrate with enough force to induce crushing/cracking in the microbeads, and indicate operational success of our piezo-actuated silicon impactor designs within the integrated silicon chip/ microfluidic device environment as proof of concept. A caveat: the crushing efficiency of the bSi nano-needle chip may be an overestimate because we cannot rule out breakage of the fine bSi nano-needle structures during piezo-actuated crushing; fragments of bSi nano-needles in the effluent may distort our estimates of microbead fragments.

### C. Device Actuation on Yeast Cells

We also tested micromechanical impaction to crush *S. cerevisiae* using our setup. *S. cerevisiae* is a species of yeast commonly used to study fundamental biological mechanisms [22], synthesize biologics [23] etc. These yeast cells are ellipsoidal [24] in shape, typically range from ∼3-6 µm in size [25] and have cell walls ranging from 100-200 nm in thickness [25]. Even though the KOH pyramids appeared to have highest efficiency in crushing microbeads, we reasoned that the spacing between pyramids is large enough for the *S. cerevisiae* cells to escape into; hence we chose the second most efficient silicon impactor chip, i.e., the ridge geometry for these experiments. We also observed that the ridge geometry is mechanically robust and less prone to damage during fabrication and handling.

SEM imaging was performed on cell lysate collected as effluent (**Fig. 5a-d**). Though *S. cerevisiae* cells match the porous silica microbeads in size, they do not fragment into multiple pieces, unlike the (more brittle) silica beads. Instead, the cells show a flattened morphology [26] with tears on the cell walls (yellow arrow, **Fig. 5a, b**) or a doughnut morphology (yellow arrow, **Fig. 5c, d**) after undergoing impaction. *S. cerevisiae* cells are not amorphous solid spheres like the silica beads, but, with rigid cell walls and aqueous contents, they resemble hollow spheres with rigid shells instead. We posit that, under impaction, *S. cerevisiae* cell walls may collapse or develop a dent (like a dented ping pong ball), causing the donut morphology.

**Fig. 5.**
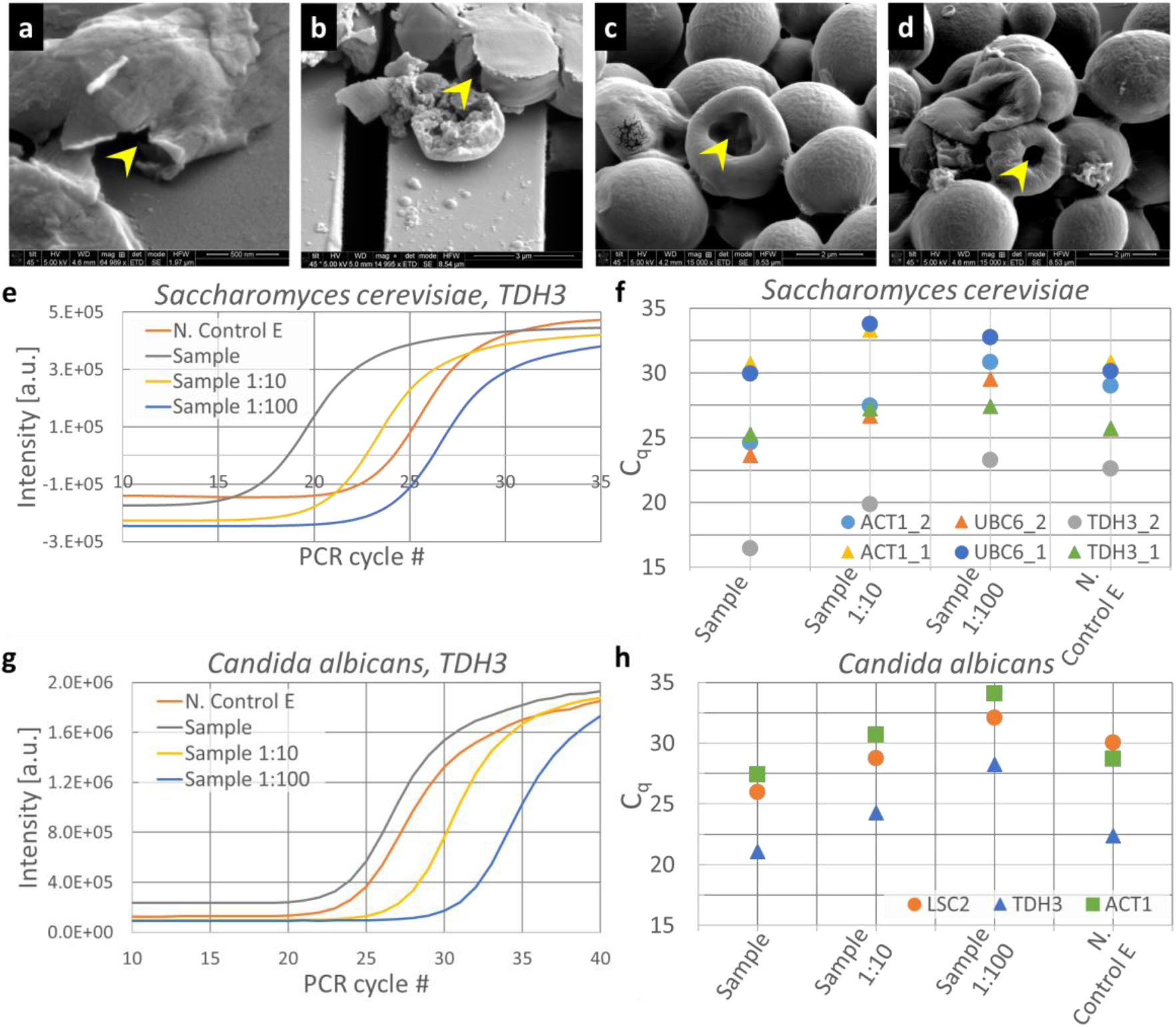
Using the silicon chip embedded microfluidic device to crush *S. cerevisiae* and *C. albicans* cells for lysis and release of mRNA. (a-d) SEM images of *S. cerevisiae* crushed using a device with ridged silicon chip. (e-h) qPCR amplification curves for *S. cerevisiae and C. albicans* lysates collected after the cells were crushed with ridged silicon chip. The crushed effluent, ‘Sample’ was tested at stock concentration, 1:10 (‘Sample 1:10’) and 1:100 (‘Sample 1:100’) dilutions. Effluent collected without piezo activity was used as negative control, (‘N. Control E’). mRNA for three commonly expressed genes were tested for each species: (e, f) *S. cerevisiae*: *ACT1, UBC6, TDH3*, and (g, h) *C. albicans*: *LSC2, TDH3, ACT1*. (f) C_q_ values extracted from the amplification curves for *ACT1, UBC6* and *TDH3* mRNA from crushed *S. cerevisiae* are plotted from 2 different experiments, indicated as ‘_1’ and ‘_2’. (g) PCR amplification curves for crushed C. albicans lysate, ‘Sample’ and dilutions at 10x (‘Sample 1:10’) and 100x (‘Sample 1:100’) are shown, along with uncrushed effluent (N. Control E). (h) C_q_ values extracted from the amplification curves for *LSC2, TDH3* and *ACT1* mRNA from crushed *C. albicans* lysate collected from an impaction experiment are shown for the sample at different dilutions, along with uncrushed effluent.

**Fig. 5e** shows the PCR amplification curves for *ACT1* cDNA in the effluent after impaction activity at stock concentration (marked as ‘Sample’), 10x and 100x dilutions (marked as ‘Sample 1:10’ and ‘Sample 1:100’, respectively), and uncrushed effluent (marked as ‘N. Control E’). Quantification cycle (C_q_) values were extracted from the amplification curves for these conditions for *ACT1, TDH3* and *UBC6* [19] (**Fig. 5f**). As the gene with the highest mRNA expression in *S. cerevisiae* [21], we expect *TDH3* to have the lowest C_q_ value of the three genes tested. The crushed cell samples have C_q_ values for *TDH3* ranging from 16.4 in experiment 1 to 25.3 in experiment 2. We suspect this is due to sedimentation of cells within the device, which could cause slightly different concentrations of cells between experiments. In each case however, the C_q_ of the crushed sample was lower than the C_q_ of the corresponding uncrushed sample control. When compared to uncrushed effluent (N. Control E), the crushed cells have a lower mean C_q_ (ΔC_q_ = 3.3), indicating that the silicon chip impaction indeed caused a fraction of the *S. cerevisiae* cell walls to lyse and release mRNA into the effluent. This holds for *ACT1* and *UBC6* also, with mean ΔC_q_ values of 2.2 and 1.1, respectively.

Compared to the *S. cerevisiae* positive control (P.C.), where 50 million *S. cerevisiae* cells were lysed and eluted in 100 µL, the crushed effluent has a concentration of 50 million cells/mL. C_q_ values of *S. cerevisiae* P.C. at different dilutions are shown in **Supplementary Fig. S3a**. If we assume 100% lysis in both Sample and P.C., the C_q_ value of the Sample should be comparable to P.C. at 1:10 dilution; a C_q_ comparable to P.C. at 1:20 dilution would indicate around 50% lysis, P.C. at 1:100 dilution around 10% lysis, etc. C_q_ values of crushed *S. cerevisiae* lysate typically lay between P.C. 1:100 and P.C. 1:1000, indicating that our lysis efficiency is <10%.

We also tested *C. albicans* cells at 50 million cells/mL for lysis using our setup. As before, qRT-PCR was used to detect the presence of *C. albicans* mRNA in the crushed lysate that was collected as effluent. Amplification curves for *TDH3* cDNA in crushed (‘Sample’) and uncrushed effluents are shown as function of PCR cycles (**Fig. 5g**), along with 10x and 100x dilutions of the crushed sample. C_q_ values extracted from the amplification curves of *ACT1, LSC2* and *TDH3* cDNA (**Fig. 5h**) show a small but consistent difference in C_q_ between crushed and uncrushed effluents. *TDH3*, being one of the top expressed genes in *C albicans*, is detected at a lower PCR cycle number (21.1) compared to *LSC2* and *ACT1* (25.9 and 27.4, respectively).

However, the cDNA for these genes are consistently detected at higher PCR cycle numbers, compared to *S. cerevisiae*, across multiple experiments (data not shown). *C. albicans* possess thicker cell walls than *S. cerevisiae* [27] and is therefore more difficult to lyse. We posit that we have a lower efficiency in crushing *C. albicans* cells on our setup compared to *S. cerevisiae*, resulting in lower numbers of mRNA in the crushed lysates.

As before, 50 million *C. albicans* cells were lysed with zymolyase and eluted in 100 µL at stock concentration and serial dilutions were used as positive controls (‘P.C.’) to estimate lysis efficiency of *C. albicans* cells on our setup (**Supplementary Fig. S3b**). Comparison of C_q_ values from Sample and P.C. estimates lysis efficiency <10% for *C. albicans*, using our setup.

We acknowledge two drawbacks in our experimental setup currently:

#### Low efficiency

Efficiency in crushing beads or microbes is low in our setup. When beads do make successful contact with the silicon chip features and the glass surface, crushing is efficient. However, not all beads that flow through the device are guaranteed to make contact with the silicon impactor chip. There are two possible reasons behind this: 1) poor bonding PDMS and glass around the silicon chip, leading to fewer beads flowing between the glass substrate and patterned silicon microarray than intended, and 2) the actuation process does not accommodate dead volume in the channel below it (i.e., when the silicon chip is pressing down, the displaced volume of fluid, carrying additional beads, squeezes out of area covered by the chip in the channel). When the chip is raised, the fluid volume rushes back under it, but the overall flow in the device also pushes the total fluid volume forward towards the outlet, carrying some beads that may never have made contact with the silicon chip. This may be addressed in the future by modifying the integrated package design to accommodate (by capturing or recirculating) the fluid escaping silicon chip impaction.

#### Inconsistency between experiments

We note some variation in bead or cell crushing efficiency in our setup. This may be due to variations in cell or bead density in the syringe and connection tubing due to sedimentation. Matching the density of the fluid with the bead or cell densities will help mitigate this issue.

All experiments were performed at a fixed flow rate, piezo amplitude, frequency, and duty cycle; these experimental parameters were not varied systematically to optimize bead-crushing or cell lysis efficiency. These optimizations are planned in future.

## IV. SUMMARY AND CONCLUSIONS

We present a packaged device and integration technique that incorporates semiconductor-based components like silicon microstructures and functionality into soft microfluidics for the lysis of microbial cells for genomics applications. We describe lithography techniques for fabricating different microstructures on silicon chip, including pyramids, pillars, ridges, dense pointed structures (cryo pyramids) and nano-needles, all of which may be used to break live microbial cell walls through micromechanical impaction to perform cell lysis. We describe the packaging of silicon chip into 3D printed microfluidic devices and the operation of the integrated device using externally coupled piezo-electric transducer and syringe pump under optical imaging. Synthetic microbeads (polystyrene, silica) of different sizes are tested as proxy for microbial cells of similar shape and size; the microbead fragments are quantified using SEM to estimate crushing efficiency that ranged between 3.7-50.4%, depending on bead size and silicon chip geometry. Two microbial species, *S. cerevisiae* and *C. albicans*, are tested for lysis by micromechanical impaction using our setup. We used qRT-PCR in addition to SEM to verify lysis of microbial cells using our setup, and estimate lysis <10 % for both species.

Considering the size of many cells of interest, future experiments will focus on optimizing piezo actuation and flow parameters using silicon chips with ridge and nano-needle geometries. Further, more active or passive components may also be integrated into the system for on-chip characterization of microbial lysate, including imaging, spectroscopy or chemical assays.

## ACKNOWLEDGMENTS

We thank Dylan Cook for his helpful comments on the manuscript. We thank Christina S. Miller, Liliana Stan and Trevor Wood for help with the dicing saw, e-beam evaporator, sputtering and qPCR help and the CNM Nanofabrication group for providing excellent work conditions in the Clean Room.

This work was supported by the University of Chicago BSD FY22 Pilot Projects funding, NIH DP2AI158157 (A.B.), Vannevar Bush Fellowship (S.G.) under the program sponsored by the Office of the Undersecretary of Defense for Research and Engineering (OUSD (R&E)) and The Office of Naval Research as the executive manager for the grant. E.R.L. was supported by the Quad Undergraduate Research Scholars Program.

Worked performed at the Center for Nanoscale Materials, a U.S. Department of Energy Office of Science user facility, was supported by the U.S. DOE, Office of Basic Energy Sciences, under Contract No. DE-AC02-06CH11357.

## DATA AVAILABILITY

The data that supports the findings of this study are available within the article.

## V. FIGURES

**Supplementary Fig. S1.**
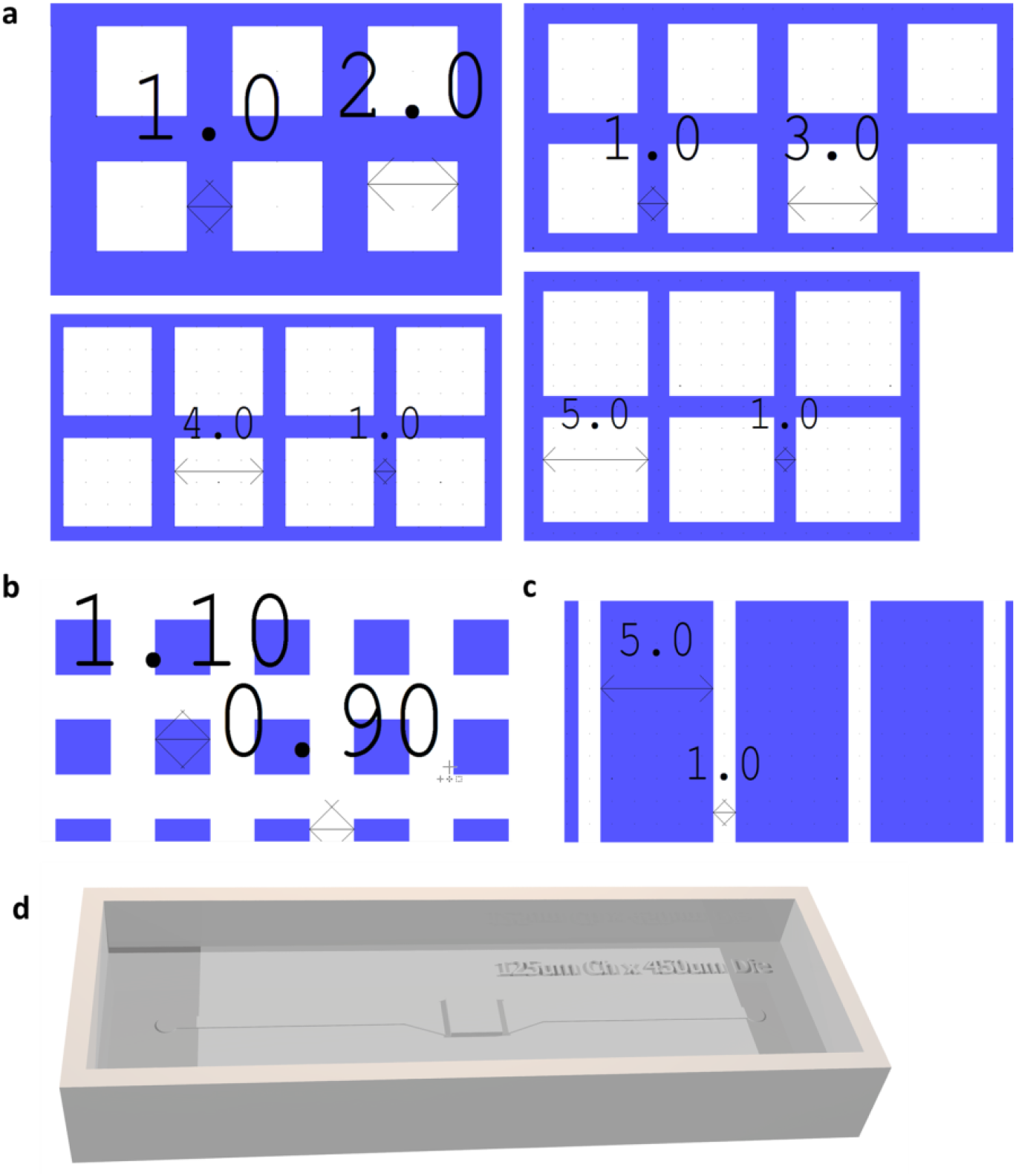
Summary of designs used for fabrication. (a) Four different layouts of KOH pyramids; (b) Pillars and Cryo pyramids; (c) Ridges; and (d) 3D printed mold to cast PDMS device.

**Supplementary Fig. S2.**
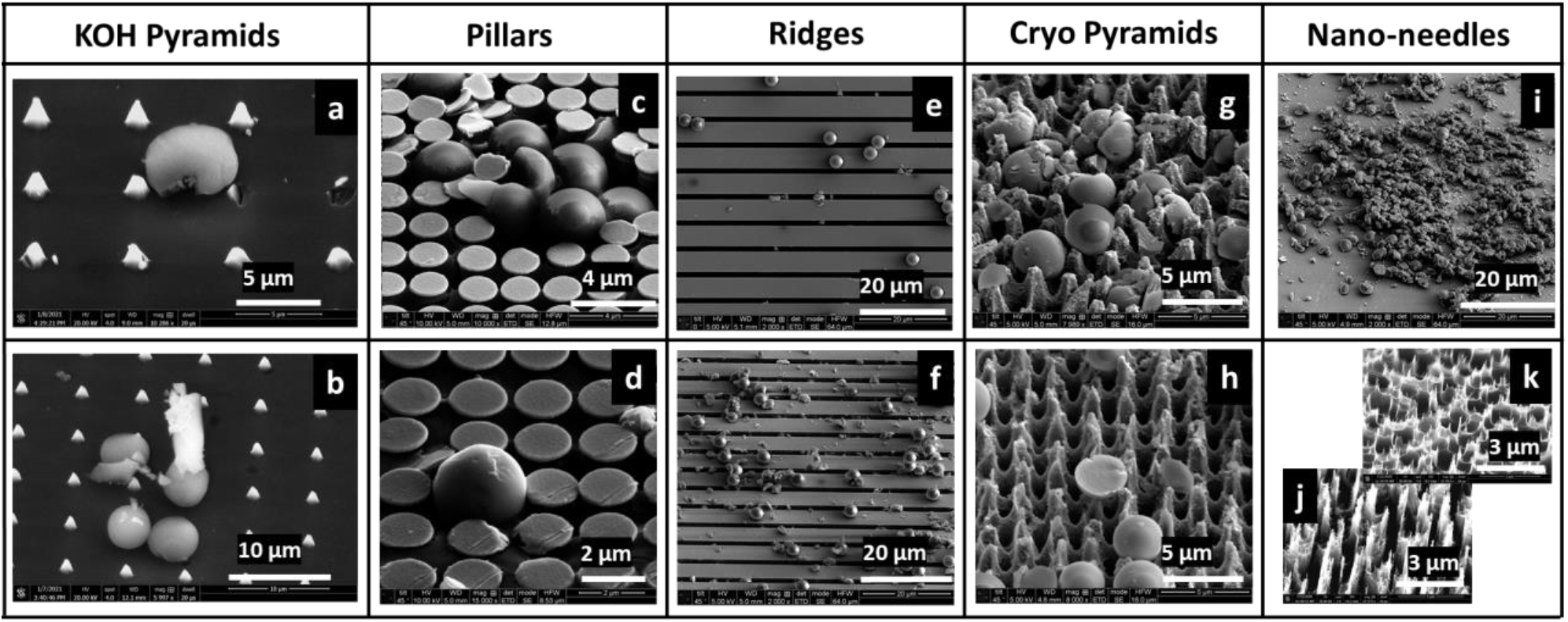
SEM images show (a-b) beads being crushed or ripped off with broken Si structures embedded into them, (c-d) beads, crushed or uncrushed, that are trapped between cylindrical pillars, (e-f) crushed beads trapped between ridges; fewer crushed beads are trapped when the ridges are perpendicular (f) to the direction of the flow compared to when with parallel with the direction of flow; (g-h) beads damaged by dense cryo pyramids, (i) nano-needles fabricated for 5 min etch, (j) relatively fragile nano-needles are generates when etched for 8 min.

**Supplementary Fig. S3.**
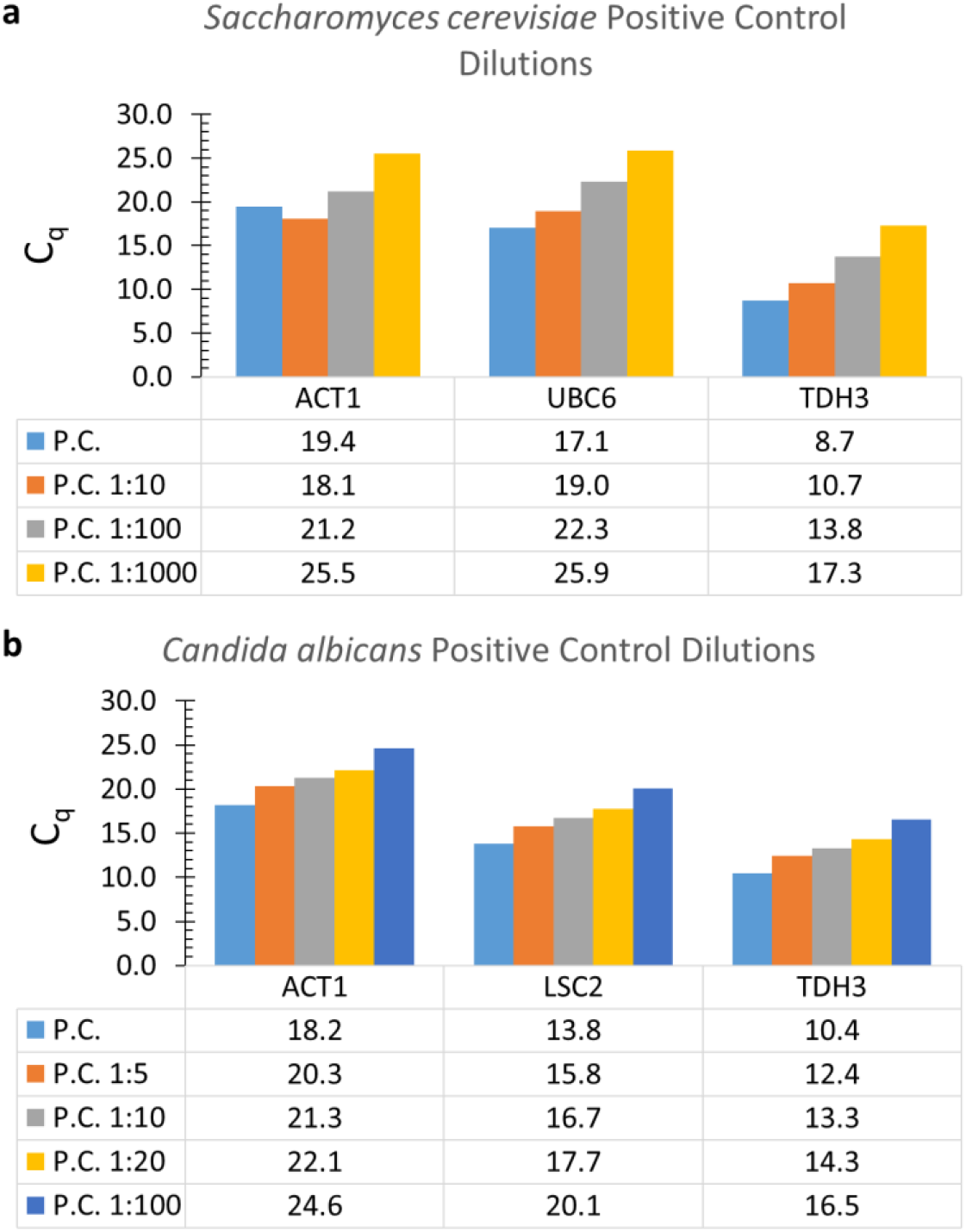
C_q_ values of (a) *S. cerevisiae* and (b) *C. albicans* lysates used as positive controls (P.C.). In each case, 50 million cells were treated with zymolyase and sarkosyl for lysis. The lysates were purified and eluted in 100 µL buffer, and different dilutions of the lysate were amplified using ACT1, UBC6 and TDH3 primers for *S. cerevisiae* and ACT1, LSC2 and TDH3 primers for *C. albicans* that were designed for each microbial species.

## REFERENCES

1. Whitesides GM. The origins and the future of microfluidics. Nature. 2006;442(7101):368–73. doi: 10.1038/nature05058.

2. Thompson AM, Paguirigan AL, Kreutz JE, Radich JP, Chiu DT. Microfluidics for single-cell genetic analysis. Lab Chip. 2014;14(17):3135–42. doi: 10.1039/c4lc00175c. PubMed PMID: 24789374.

3. Jariani A, Vermeersch L, Cerulus B, Perez-Samper G, Voordeckers K, Van Brussel T, et al. A new protocol for single-cell RNA-seq reveals stochastic gene expression during lag phase in budding yeast. eLife. 2020;9:e55320. doi: 10.7554/eLife.55320.

4. Urbonaite G, Lee JTH, Liu P, Parada GE, Hemberg M, Acar M. A yeast-optimized single-cell transcriptomics platform elucidates how mycophenolic acid and guanine alter global mRNA levels. Communications Biology. 2021;4(1):822. doi: 10.1038/s42003-021-02320-w.

5. Shehadul Islam M, Aryasomayajula A, Selvaganapathy PR. A Review on Macroscale and Microscale Cell Lysis Methods. Micromachines (Basel). 2017;8(3):83. doi: 10.3390/mi8030083. PubMed PMID: PMC6190294.

6. Carlo DD, Jeong K-H, Lee LP. Reagentless mechanical cell lysis by nanoscale barbs in microchannels for sample preparation. Lab Chip. 2003;3(4):287–91. doi: 10.1039/B305162E.

7. Kobayashi-Kirschvink KJ, Gaddam S, James-Sorenson T, Grody E, Ounadjela JR, Ge B, et al. Raman2RNA: Live-cell label-free prediction of single-cell RNA expression profiles by Raman microscopy. bioRxiv. 2022:2021.11.30.470655. doi: 10.1101/2021.11.30.470655.

8. Choi C-H, Kim C-J. Fabrication of a dense array of tall nanostructures over a large sample area with sidewall profile and tip sharpness control. Nanotechnology. 2006;17(21):5326–33. doi: 10.1088/0957-4484/17/21/007.

9. Ribbing C, Cederstrom Br, Lundqvist M. Microfabrication of saw-tooth refractive x-ray lenses in low-Z materials. Journal of Micromechanics and Microengineering. 2003;13(5):714–20. doi: 10.1088/0960-1317/13/5/325.

10. Seidel H, Csepregi L, Heuberger A, Baumgärtel H. Anisotropic Etching of Crystalline Silicon in Alkaline Solutions: I. Orientation Dependence and Behavior of Passivation Layers. Journal of The Electrochemical Society. 1990;137(11):3612–26. doi: 10.1149/1.2086277.

11. Belougne J, Ozerov I, Caillard C, Bedu F, Ewbank JJ. Fabrication of sharp silicon arrays to wound Caenorhabditis elegans. Scientific Reports. 2020;10(1):3581. doi: 10.1038/s41598-020-60333-7.

12. Zubel I, Kramkowska M. The effect of isopropyl alcohol on etching rate and roughness of (1 0 0) Si surface etched in KOH and TMAH solutions. Sensors and Actuators A: Physical. 2001;93(2):138–47. doi: https://doi.org/10.1016/S0924-4247(01)00648-3.

13. Boer MJd, Gardeniers JGE, Jansen HV, Smulders E, Gilde MJ, Roelofs G, et al. Guidelines for etching silicon MEMS structures using fluorine high-density plasmas at cryogenic temperatures. Journal of Microelectromechanical Systems. 2002;11(4):385–401. doi: 10.1109/JMEMS.2002.800928.

14. Kamto A, Divan R, Sumant AV, Burkett SL. Cryogenic inductively coupled plasma etching for fabrication of tapered through-silicon vias. Journal of Vacuum Science & Technology A. 2010;28(4):719–25. doi: 10.1116/1.3281005.

15. Michalska M, Gambacorta F, Divan R, Aranson IS, Sokolov A, Noirot P, et al. Tuning antimicrobial properties of biomimetic nanopatterned surfaces. Nanoscale. 2018;10(14):6639–50. doi: 10.1039/C8NR00439K.

16. Ivanova EP, Hasan J, Webb HK, Gervinskas G, Juodkazis S, Truong VK, et al. Bactericidal activity of black silicon. Nature Communications. 2013;4(1):2838. doi: 10.1038/ncomms3838.

17. McDonald JC, Duffy DC, Anderson JR, Chiu DT, Wu H, Schueller OJA, et al. Fabrication of microfluidic systems in poly(dimethylsiloxane). ELECTROPHORESIS. 2000;21(1):27–40. doi: https://doi.org/10.1002/(SICI)1522-2683(20000101)21:1<27::AID-ELPS27>3.0.CO;2-C.

18. Shehadat SA, Gorduysus MO, Hamid SSA, Abdullah NA, Samsudin AR, Ahmad A. Optimization of scanning electron microscope technique for amniotic membrane investigation: A preliminary study. Eur J Dent. 2018;12(4):574-8. Epub 2018/10/30. doi: 10.4103/ejd.ejd_401_17. PubMed PMID: 30369805; PubMed Central PMCID: PMCPMC6178683.

19. Teste MA, Duquenne M, Francois JM, Parrou JL. Validation of reference genes for quantitative expression analysis by real-time RT-PCR in Saccharomyces cerevisiae. BMC Mol Biol. 2009;10(1471-2199 (Electronic)):99. Epub 2009/10/31. doi: 10.1186/1471-2199-10-99. PubMed PMID: 19874630; PubMed Central PMCID: PMCPMC2776018.

20. Nailis H, Coenye T, Van Nieuwerburgh F, Deforce D, Nelis HJ. Development and evaluation of different normalization strategies for gene expression studies in Candida albicans biofilms by real-time PCR. BMC Molecular Biology. 2006;7(1):25. doi: 10.1186/1471-2199-7-25.

21. Dohn R, Xie B, Back R, Selewa A, Eckart H, Rao RP, et al. mDrop-Seq: Massively Parallel Single-Cell RNA-Seq of Saccharomyces cerevisiae and Candida albicans. Vaccines. 2022;10(1). doi: 10.3390/vaccines10010030.

22. Altmann K, Dürr M, Westermann B. Saccharomyces cerevisiae as a Model Organism to Study Mitochondrial Biology. In: Leister D, Herrmann JM, editors. Mitochondria: Practical Protocols. Totowa, NJ: Humana Press; 2007. p. 81–90.

23. Walker RSKAUPISTIAoYSBGttPoB. Genes [Internet]. 2018; 9(7).

24. Saito TL, Ohtani M, Sawai H, Sano F, Saka A, Watanabe D, et al. SCMD: Saccharomyces cerevisiae Morphological Database. Nucleic Acids Res. 2004;32(Database issue):D319-22. Epub 2003/12/19. doi: 10.1093/nar/gkh113. PubMed PMID: 14681423; PubMed Central PMCID: PMCPMC308847.

25. Ahmad MR, Nakajima M, Kojima S, Homma M, Fukuda T. The effects of cell sizes, environmental conditions, and growth phases on the strength of individual W303 yeast cells inside ESEM. IEEE Trans Nanobioscience. 2008;7(3):185-93. Epub 2008/09/10. doi: 10.1109/TNB.2008.2002281. PubMed PMID: 18779098.

26. Nowlin K, Boseman A, Covell A, LaJeunesse D. Adhesion-dependent rupturing of Saccharomyces cerevisiae on biological antimicrobial nanostructured surfaces. Journal of The Royal Society Interface. 2015;12(102):20140999. doi: 10.1098/rsif.2014.0999.

27. Klis Frans M, de Koster Chris G, Brul S. Cell Wall-Related Bionumbers and Bioestimates of Saccharomyces cerevisiae and Candida albicans. Eukaryotic Cell. 2014;13(1):2–9. doi: 10.1128/EC.00250-13.

